# Comprehensive re-analysis of hairpin RNAs in fungi reveals ancestral links

**DOI:** 10.1101/2022.09.15.508153

**Authors:** Nathan R. Johnson, Luis F. Larrondo, José M. Álvarez, Elena A. Vidal

## Abstract

RNA interference is an ancient mechanism with many regulatory roles in eukaryotic genomes, with small RNAs acting as their functional element. While there is a wide array of classes of small-RNA-producing loci, those resulting from stem-loop structures (hairpins) have received profuse attention. Such is the case of microRNAs (miRNAs), which have distinct roles in plants and animals. Fungi also produce small RNAs, and several publications have identified miRNAs and miRNA-like (mi/milRNA) hairpin RNAs in diverse fungal species using deep sequencing technologies. Despite this relevant source of information, relatively little is known about mi/milRNA-like features in fungi, mostly due to a lack of established criteria for their annotation. To systematically assess mi/miRNA-like characteristics and annotation confidence, we searched for publications describing mi/milRNA loci and re-assessed the annotations for 40 fungal species. We extracted and normalized the annotation data for 1,677 reported mi/milRNA-like loci and determined their abundance profiles, concluding that less than half of the reported loci passed basic standards used for hairpin RNA discovery. We found that fungal mi/milRNA are generally more similar in size to animal miRNAs and were frequently associated with protein-coding genes. The compiled genomic analyses identified 18 mi/milRNA loci conserved in multiple species. Our pipeline allowed us to build a general hierarchy of locus quality, identifying around 200 loci with high-quality annotations. We provide a centralized annotation of identified mi/milRNA hairpin RNAs in fungi which will serve as a resource for future research and advance in understanding the characteristics and functions of mi/milRNAs in fungal organisms.

## Introduction

Silencing by RNA interference (RNAi) is an ancient system for regulating RNA abundance found within eukaryotes. Small regulatory RNAs (sRNAs) are the functional elements behind RNAi and are typically 20-24 nucleotides in length. Functionally, sRNAs play key roles in genome stability [1,2], protection against RNA-based organisms [3,4], and regulation of gene expression [5]. In the context of pathogenic organisms, such as fungal plant pathogens, sRNAs have even been shown to display *trans*-kingdom functions [6,7], where fungal sRNAs serve as effectors of pathogenicity, regulating host genes to undermine resistance.

The biogenesis pathway of a sRNA locus can provide key insights into its function in an organism. Significantly, pathways differ by the sRNA-class and by organism but generally follow the same schema [8,9]. Regions of double-stranded RNA (dsRNA) are processed by a Type-III RNAse called Dicer or Dicer-like (collectively abbreviated here as DCR), producing sRNA duplexes which are subsequently loaded in an Argonaute protein (AGO) for their resulting function. DCRs are responsible for the resulting sRNA length [10] and these lengths are selective for the specific AGO loading. As a result, DCRs directly influence sRNA function [11]. The source of the dsRNA is also critical to sRNA function. Two pathways exist for this: 1) dsRNA produced through synthesis by an RNA-dependent RNA polymerase (RDR), or 2) complementary regions in transcripts form stem-loop fold-back structures known as hairpins [12]. Several RDR-derived sRNA classes have been defined in eukaryotes, with fungal types including those resulting from DNA damage (qiRNAs) [13], associated with meiotic silencing [14], or associated with transcriptional silencing (i.e. quelling) [15]. Hairpin-derived sRNAs (hpRNAs) are widespread in eukaryotes and include microRNAs (miRNAs), which have central roles in gene expression control in plants and animals.

Highly-precise dicing characterizes miRNAs, with the most-abundant sequences (MASs) coming from a single duplex in the 5’ and 3’ hairpin arms [16]. These have been sometimes referred to as the mature-miRNA or the miR and the miR*, though these terms are imprecise considering both of these sequences may be functional and may reverse in the rank of abundance [17,18]. Homologs of miRNAs are classified into families, defined by nearly identical mature-miRNA sequences, though more distant relationships have been shown to exist [19]. The mechanistic function of miRNAs is distinctly different by clade. In plants, miRNAs function through direct cleavage of a target mRNA [20], whereas in animals the process is less straight-forward, relying on inhibition of translation and mRNA de-tailing, de-capping, and degradation by exonucleases [21]. A significant number of miRNA are anciently conserved in sequence and function [22], though they may also be highly clade- or species-specific [23,24]. There are also examples of hpRNAs that are clearly divergent from miRNAs and produce a spectrum of sizes from differing pathways [12]. This phenomenon exists natively in plants [25,26] and has long been a biotechnological tool for induction of RNAi [27], however, further exploration is needed to define and confirm these loci in more organisms.

Fungi have also been shown to produce hpRNAs. The hpRNA class of miRNA-like RNAs (milRNAs) was first identified in *Neurospora crassa*, where they follow the same schema: RNA foldbacks are processed by one of two redundant DCRs (NcDCL1 and NcDCL2), which are subsequently loaded into an AGO protein (QDE2) [28]. Homolog proteins of the DCR and AGO families have been identified in many fungal species [29] and are involved in milRNA biogenesis and function. Since, over 60 publications have cited hpRNAs in fungi, using the designations milRNA or miRNA (collectively referred to as mi/milRNAs). In fungi, these sRNAs have been identified frequently in the context of pathogenesis [30]. This also includes some proposed to regulate gene expression in *trans* [31–33], regulating plant and animal host gene expression. However, most lack clear evidence for the causative role of the sRNA in the process, as many studies rely on target prediction alone for understanding their biological role.

The advent of small-RNA-seq technology has massively increased the amount of data available for detecting sRNA-producing loci. This has affected the standards of miRNA identification, as we now can rely on high-quality evidence for their identification, as opposed to low-throughput and error-prone approaches like identification through RT-PCR. A wide spectrum of tools are available for annotation of miRNAs, with most of them based on strict rules derived from characteristics that are specific to known miRNAs from plants or animals [34–37]. Despite many publications identifying mi/milRNA loci in fungi, there has yet to be a systematic assessment of their characteristics and annotation confidence, as in other eukaryotic species [16,38]. In plants and animals, much is known about miRNAs. Here, we know the lengths of the hairpins and their foldback dynamics, the sizes of sRNAs that are produced, and the mechanisms behind their targeting [16,39]. We also know the genomic context for the hairpins and the extent of their evolutionary conservation. In fungi we lack this categorical knowledge about mi/milRNAs, which is further complicated by the abundance of non-uniform pipelines and inconsistent methodology, raising significant questions about the quality of their systematic identification. In this work, we focused solely on sRNAs derived from hairpin structures, aiming to identify fundamental characteristics of fungal mi/milRNAs. Furthermore, we sought to assess the quality of their annotations based on key metrics, ultimately building a centralized and well-documented annotation for public use. In this effort, we provide the first steps to evaluate the actual shape, size, context, and function of mi/milRNAs in fungi.

## Results

### Annotation of mi/milRNA in fungal species is sparse and heterogeneous

To gain insights into fungal mi/milRNA distinctive features and to assess the state of the genomic annotation of published loci, we conducted an exhaustive search for publications including terms related to fungal miRNAs and milRNAs in PubMed. The search yielded a list of 58 publications which assess fungal mi/milRNAs, 51 of which performed and reported *de novo* mi/milRNA loci identification from sRNA-seq data (Figure 1A, Table S1), using a broad range of bioinformatic tools or pipelines (Figure 1A). Noteworthy, these tools have been developed for miRNA prediction and annotation in plants and/or animals and can fail to identify loci with particular characteristics of mi/milRNA in fungi, as we demonstrated below. The majority of these sRNA-seq libraries are publicly available (Figure 1A, Table S1) and span 40 fungal species, including some subspecies (Figure 1B), presenting a prime opportunity to uncover common fungal mi/milRNA characteristics. These species come from a variety of lifestyles and relevance to humans, including laboratory models as well as many plant and animal pathogens (Table S2). Nearly all explored species are Ascomycetes and Basidiomycetes, though more distant divisions of Glomeromycota and Microsporidia are also included (Figure 1B, Table S2). Several species have many observed mi/milRNA loci, such as *Sclerotinia sclerotiorum* (*Scscl*) or *Fusarium graminearum* (*Fugra*) (Figure 1B). However, these counts may be more related to the study or the particular number of studies for a given organism (Figure S1). This could be due to differing levels of bioinformatic and/or experimental evidence required for mi/milRNA annotation and may fail to reveal their actual abundance/diversity in particular fungal species. For instance, only around half of the publications provide coordinates or complete sequences for precursor genes (Figure 1A), key information to derive structural and compositional features of the hairpins and genomic contexts. Furthermore, less than half of the published mi/milRNA have support from at least two independent biological replicates (Figure 1A), a requirement regarded as vital for miRNA annotation [37]. Regarding experimental validation, a minority of the source publications for a locus provide genetic evidence to support mi/milRNA biogenesis machinery using knock-out or over-expressor lines, or explore targeting relationships with direct experimental evidence (Figure 1A). This assessment indicates a need for establishing a set of standard rules for mi/milRNA annotation in fungi, as it has been described for animals and plants [16,37].

**Figure 1.**
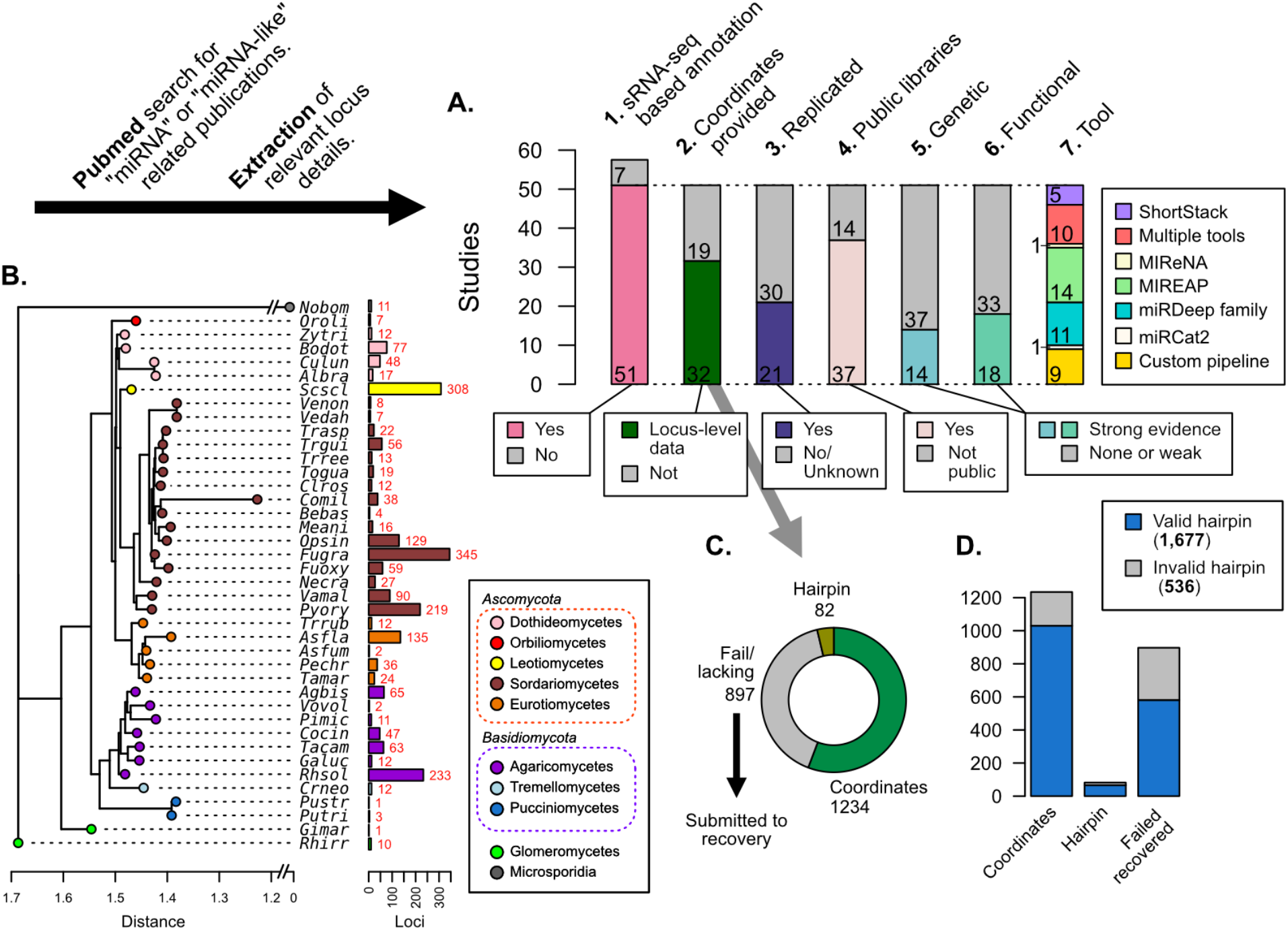
Publications referring to miRNAs and/or miRNA-like RNAs in fungi were identified, classified, and assessed for reported loci. **A)** Breakdown of important metrics in fungal mi/milRNA loci identified in publications (Table S1). 1. Number of published studies reporting sRNA-seq-based annotations. 2. Number of publications where genomic coordinates are given for each mi/milRNA precursor gene. 3. Number of studies that have 2 or more library replicates for a condition. 4. Number of studies that make the sRNA-seq data available through a public repository. 5. Number of studies that support the synthesis pathway for the mi/milRNA through knock-out or over-expression lines. 6. Number of studies that give evidence for the function of a mi/milRNA through target-site manipulation or molecular evidence of cleavage (evidence quality designations described in Methods). 7. Number of studies using a given tool/pipeline to discover mi/milRNA loci. **B)** Dendrogram constructed from 18S rRNA sequences for the species with reported loci in A1 (MUSCLE alignment, RAxML maximum likelihood tree). Colored tips represent the taxonomic class for each species identified by SILVA. Names are given as a 5-letter abbreviation (full names in Table S2). Number of loci identified is shown by colored bars. **C)** Assessment of the quality of loci reporting for published mi/milRNA loci. Loci are classified by whether they have a reported sequence and genomic coordinates for the entire precursor hairpin (Coordinates, green), only reported sequence for the precursor hairpin without genomic coordinates (Hairpin, olive green), or have insufficient or no information regarding the hairpin locus (Fail/lacking, grey). Loci in this final category were subjected to a recovery pipeline to characterize their genomic coordinates, as explained in Methods. **D)** An assessment of hairpins relative to their source evidence as explained in C. Valid hairpins are defined as sequences containing the proposed most-abundant sequence, with no secondary structures or more than 20 unpaired bases in the duplex pairing region.

### Loci lacking coordinates were recovered using genome-based inference

Reporting sequence information and genomic coordinates of mi/milRNA loci is essential to determine precursor features. Around one third of loci had neither an annotation associated with the full precursor sequence nor complete genomic coordinates (Figure 1C). To gain insights into the genomic origin of these hairpins, we utilized bowtie alignment to identify perfect matches of a supplied mature RNA sequence to the corresponding genome version and modeled possible precursor hairpin structures with RNAfold [40].

This hairpin precursor recovery pipeline was evaluated in terms of precision and sensitivity for identifying the correct locality for a mi/milRNA locus as well as its folding pattern, confirming that our pipeline has reasonably high accuracy for predicting the correct genomic locality, based on the number of alignments for a cited MAS for a mi/milRNA (Figure S2A). We also found solid metrics for recapitulating the same hairpin when testing the pipeline against known hairpins (Figure S2B). We considered three minimal criteria for determining whether a sequence corresponds to a valid hairpin: 1) the precursor contains the reported MAS(s) within its boundaries, 2) the foldback region that gives rise to the RNA duplex must not contain secondary stems, and 3) the duplex must not contain large loops (maximum of 20 unpaired bases) [37]. Hairpins reported in publications and obtained from the recovery pipeline were evaluated according to these standards, finding that most of them met these minimum requirements (Figure 1D). Notably, around 20% of the published loci failed one or more of these tests and were excluded from further analyses.

### Independent re-assessment of reported mi/milRNA reveals inconsistent expression of expected most-abundant RNA sequences for an important proportion of loci

Current miRNA annotation pipelines depend on algorithms that mine high-throughput sRNA-seq data to determine mature miRNAs, as well as putative precursor sequences. These algorithms utilize different approaches to fulfill this task. The most commonly used tools perform or require a read alignment to genome sequences, followed by locus determination based on the genomic position and depth of the reads. Although bioinformatic prediction of sRNA loci from sequencing data is the preferred method to discover miRNAs and other types of sRNAs, frequently *bona fide* miRNA annotations require further curation and validation considering the differences in annotation standards. Hence, the quality of the annotations may appear quite variable among the submitted datasets (Figure S1) [37,38,42,43].

We sought to independently evaluate reported mi/milRNAs in each publication, taking advantage of the availability of most of the source sRNA-seq data used for annotation (Figure 1A). Libraries were obtained from public repositories and trimmed, filtered, and aligned. We confirmed alignment rates mostly similar to that of their parent publications for each step, where the data are available (Table S3). In these publications, mi/milRNA are reported with a short sequence referred to as the mature sequence or alternatively as the miRNA or milRNA. In the case of animal and plant miRNA discovery, mature miRNA sequences are the MASs aligned to a locus, as a requirement of precise locus processing [16,37]. However, within a large proportion of loci, we found that the mature sequence reported is not the MAS for the locus (Figure 2A, Table S4). Even more problematic, we found numerous examples of loci that had no reads aligning with the reported mature sequence or loci with no reads aligned at all (Figure 2A, Table S4). These categories are examples for which the locus annotation pipeline may need further curation via a consistent set of rules. Close inspection of sequences in raw, untrimmed libraries shows that it is unlikely these issues are caused by differences in the trimming, filtering or alignment steps, since the exact reported mature mi/milRNA sequence was not present in the raw libraries. In loci where another aligned sequence was found to be the MAS, this sequence was used as the corresponding mature sequence in subsequent analyses (Table S5).

**Figure 2.**
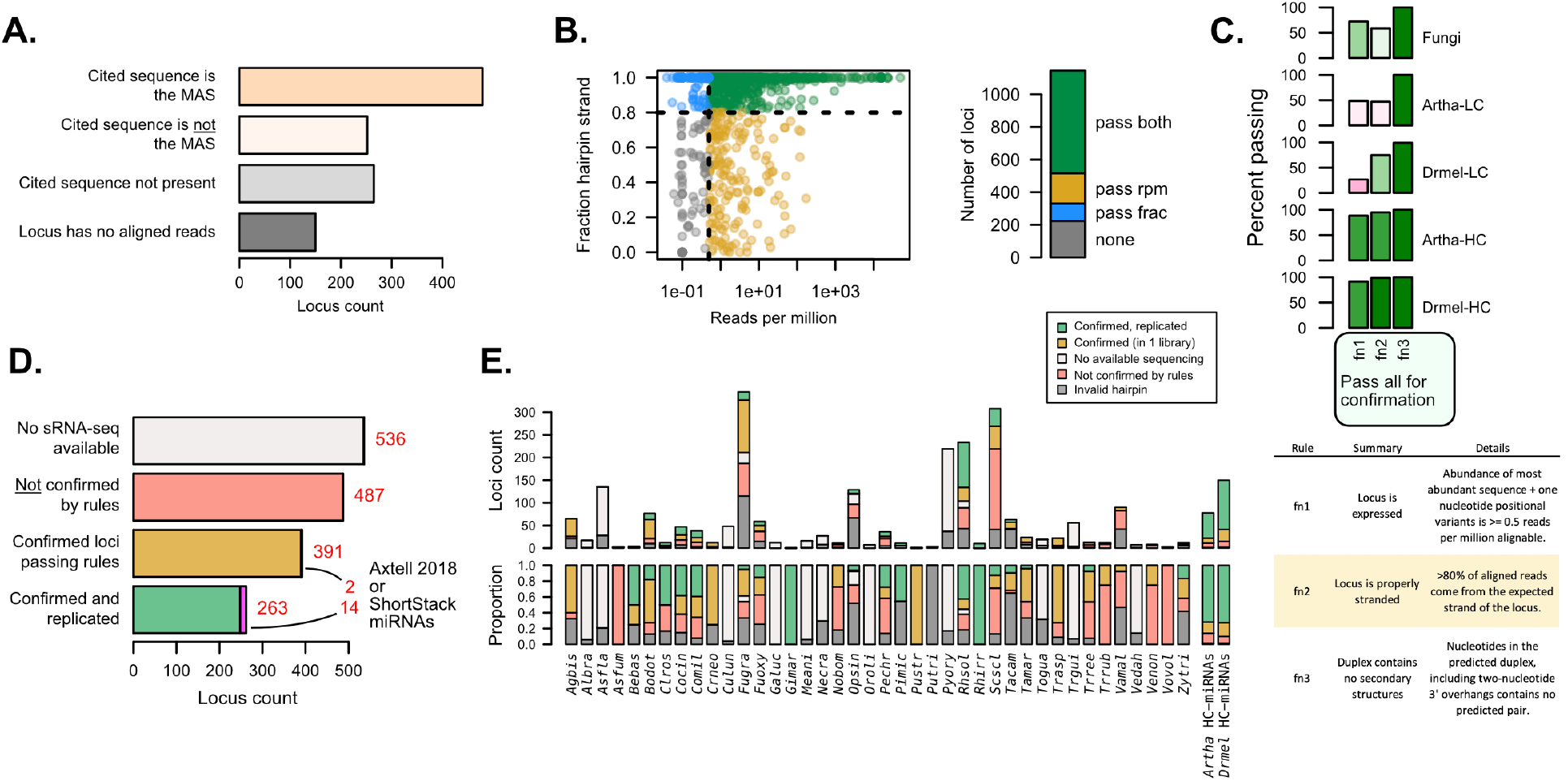
Independent evaluation of mi/milRNA annotations using sRNA-seq data. **A)** Number of hairpin loci in which the reported mature mi/milRNA sequence is the most-abundant sequence (MAS) (orange), another sequence in the locus is the MAS (pink), the cited sequence is not present in the locus (light grey), or the reported locus has no aligned reads (dark grey). **B)** Abundance (aligned reads per million) and strandedness for mi/milRNA loci. Each dot represents a locus, and the colors represent whether the locus passes the 0.5 reads per million cut-off of expression (pass rpm, yellow), has at least 80% of the aligned reads stranded (pass frac, blue), fulfills both criteria (pass both, green) or does not fulfill any of the criteria (none, grey) **C)** Proportion of loci passing the three minimal locus profile rules (fn1, fn2, and fn3, described in table below). *Arabidopsis thaliana* (*Artha*) and *Drosophila melanogaster* (*Drmel*), low (LC) and high-confidence (HC) miRNAs from miRbase tested in a single library are included as a reference. **D)** Number of loci without available source sRNA-sequence libraries (light grey), loci failing one or more rules indicated in B and C (red), loci passing all rules (yellow), and loci passing all rules in two or more independent libraries (green). For confirmed loci, we show in magenta those passing all the rules for annotating miRNAs in plants described in the ShortStack [41] and Axtell-2018 [37] rule sets. **E)** Locus count and proportion of loci as in D), separated by the parent species and including loci for which a valid hairpin could not be identified (dark grey).

### Variable rates of confirmed mi/milRNA loci among published datasets

As mentioned earlier, several sets of rules have been established for miRNA discovery and annotation in plants and animals [16,34–36]. These rules are derived from examples of known miRNA loci in well-studied organisms, and focus on the specific alignment profile of sRNA-seq reads over these loci, which is intimately related to the biogenesis mechanism of these regulatory molecules. Applying these criteria (summarized in Table S6), we found that fungal sRNAs frequently fail to meet key points compared to animal or plant miRNAs, resulting in very few that pass all rules in a set. This partial fulfillment of criteria is similar to the performance of miRNAs annotated as low confidence in miRbase for plants and animals (Figure S3). This led us to explore a less-strict set of rules to confirm that reported mi/milRNA in fungi are sRNAs derived from *bona fide* hairpins.

sRNAs originate from dsRNA precursors, either produced from an RNA-dependent polymerase or from a stem-loop fold-back in the case of hpRNAs. These scenarios result in different alignment profiles of sRNA-seq reads, as hpRNAs will only be produced from the strand of the hairpin [12]. It follows that the first basic criterion is that hpRNAs should largely originate from the proposed hairpin strand (>80%) [41] (Table S6). A second criterion focuses on a minimum standard of expression. Here, we estimated that a minimum threshold of detection for the MAS is 0.5 reads per aligned loci (Table S6) as this would be at the detection limit for libraries of depths around five million aligned reads, a threshold passed by the raw depth of ~95% of libraries in referenced in this work (Figure S4). Finally, we applied a third criterion for valid hairpins: the duplex for the MAS must contain no secondary structures (Table S6). This needed to be re-evaluated despite the prior hairpin validation (Figure 1D), due to the discovery of different MASs than reported for many loci (Figure 2A). We found that nearly half of the loci pass these three criteria in at least one library (Figure 2B, 2C).

In order to evaluate how these three rules allowed for the discrimination between high and low confidence miRNA loci, we determined the performance of known miRNA reported in miRbase for *Arabidopsis thaliana* (*Artha*) and *Drosophila melanogaster* (*Drmel*) using public sRNA-seq libraries for these two organisms. We found low-confidence miRNAs performed poorly, especially in terms of strandedness and expression, while for high-confidence loci more than 80% passed all three rules (Figure 2C). This points to the ability of these rules to discriminate low-confidence annotations. We found that fungal loci perform considerably worse than high-confidence *Artha* and *Drmel* loci in terms of passing both expression and stranding filters (Figure 2C). However, it is noteworthy that these fungal loci still perform much better than low-confidence miRNA considering these metrics (Figure 2C), indicating that they may represent a heterogeneous mixture of sRNA, possibly including siRNA or degradation-related loci, for example.

From our initial set of 1,677 loci, 390 (23.2%) passed all criteria (“confirmed” set, Figure 2D, Table S4), while 264 (15.4%) passed all criteria in two or more different libraries (“confirmed and replicated” set, Figure 2D, Table S4), for a total of 654 confirmed loci. Interestingly, 16 (0.9%) of these loci met our criteria and also the more strict set of rules used for plant miRNAs (ShortStack and Axtell-2018) [37,41]. We found important differences between the numbers of “confirmed” and “confirmed and replicated” loci between species, with some of them showing a lower number (e.g. *Trichophyton rubrum – Trrub, Verticillium dahliae* – *Vedal*) or a higher number (e.g. *Rhizophagus irregularis* – *Rhirr, Gigaspora margarita – Gimar*) of confirmed loci than *Artha* or *Drmel* (Figure 2E). Confirmation rates are highly variable in relation to source publication (Figure S5A), with lower confirmation rates tending to be associated with specific tools and pipelines (Figure S5B). While most mi/milRNA loci produce 21-24 nucleotide sRNAs as the MAS, there is no clear association of sRNA sizes with rates of confirmation (Figure S5C). In particular, species with high counts of identified loci such as *Fugra* and *Scscl* are prone to low confirmation rates pointing to systematic issues with the annotation pipelines, including problems with identifying valid hairpins (Figure 2E). This analysis further confirms the need for a standardized pipeline and set of criteria for miRNA annotation in fungi.

### mi/milRNA are frequently associated with protein coding regions

The genomic context of miRNAs is an important feature to understanding their function and evolution. In animals particularly, miRNAs can associate quite closely with protein coding genes, including originating from exons, introns, and untranslated regions [45,46]. Conversely, plant miRNAs are mostly distant from coding regions [47]. In fungi the genomic origins of mi/milRNAs have not been thoroughly studied. To gain insight into the genomic context of fungal mi/milRNAs, we first determined whether loci overlapped with any known structural or non-regulatory RNA. Structural RNAs (rRNAs, tRNAs) and non-regulatory sRNAs (snRNAs, snoRNAs) have fold patterns that might be similar to a hairpin. Considering that these are expressed sequences that likely produce short RNAs due to degradation, these may be erroneously assigned as miRNAs. Similarly, transposable elements are known to produce siRNAs [4]. For these reasons hpRNAs that overlap with other gene products should be treated with great scrutiny. We performed a blast search of all hairpin sequences against Rfam RNA families excluding RNAi-related sRNA families [48]. As shown in Figure 3A, most hairpins did not find hits in Rfam. The minority that did find hits can represent an outsized presence genomically. This appears to explain some of the most repetitive sequences such as bba-milR4 in *Beauveria bassiana* (*Bebas*), which matches the nearly universal U6 snRNA (Figure S6, Figure 3A). In this case, we find that bba-milR4 matches hundreds of loci in animals and plants (Figure S6A) and is found in the majority of other searched genomes (Figure S6B).

**Figure 3.**
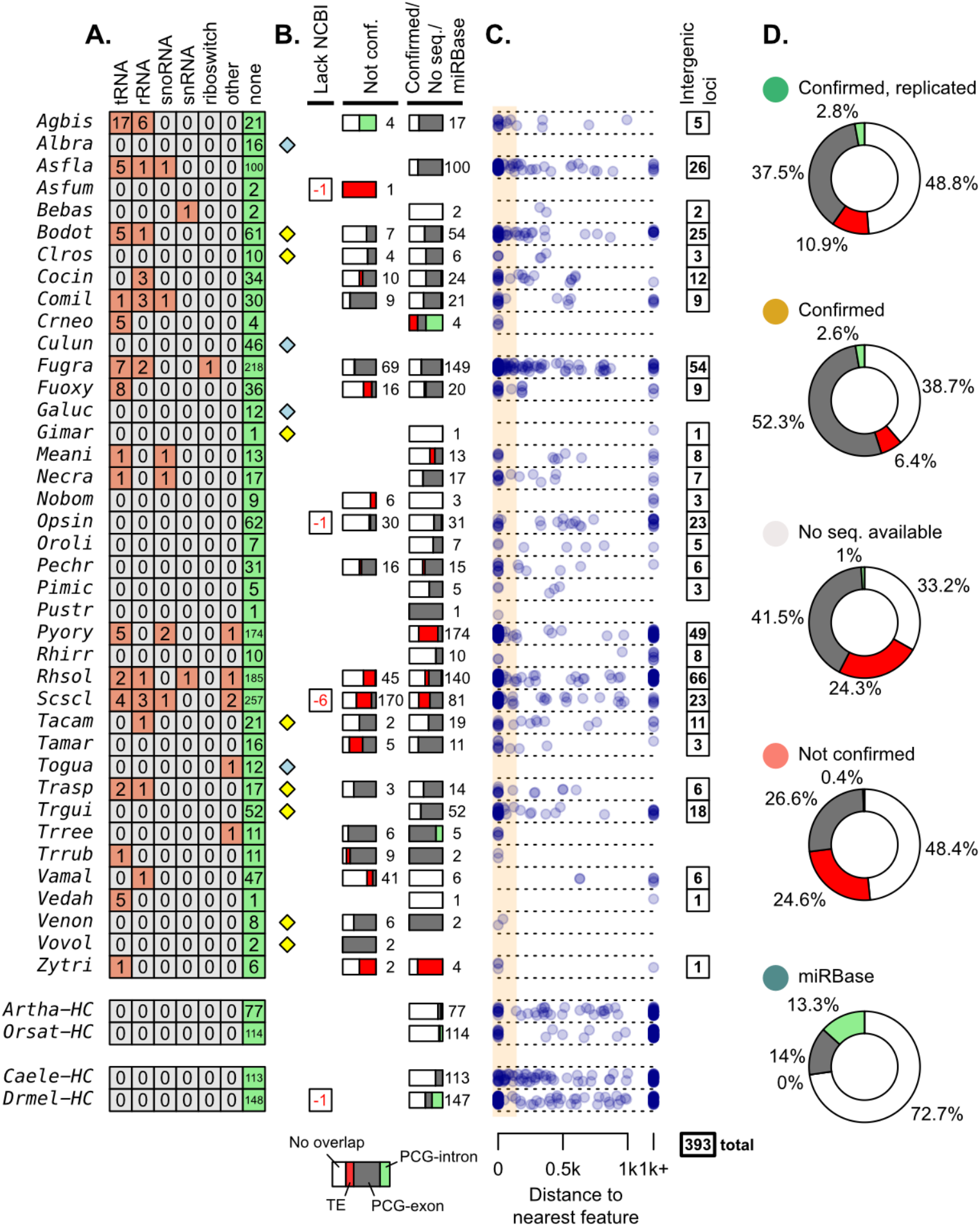
mi/milRNA loci were evaluated for homology and overlap with other genomic features. **A)** Number of loci matching Rfam structural RNA families, identified by blastn (bitscore >= 50). Categories are merged from multiple Rfam entries (Table S5). **B)** Proportions of loci originating from transposable elements (TEs, red) [44], protein-coding gene (PCG) exons (dark grey), PCG introns (green), or no overlap or feature closer than 100 nt (white). Proportion bars are composed of loci which are not found in Rfam and have a homologous sequence in an annotated genome. Bars are divided by loci by their confirmation status from Figure 2 (Not conf., not confirmed by rules; Confirmed, confirmed loci passing rules + confirmed and replicated; No seq., No sRNA-seq available). Numbers to the right of bars indicate the total locus count for a division. Diamonds indicate species which have no available TE annotations (yellow) or no TE/gene annotations (blue) available. Loci which failed to find a homologous location in the annotated NCBI assembly are shown with a negative number (red). **C)** Genomic distances to the nearest feature (nucleotides) for loci from the right bar (panel B). Those more distant than 1000 nucleotides are shown on the right. Range for considering an intersection (100 nucleotides) is shown with an orange box. Those loci which do not intersect are considered intergenic, with their counts shown on the right. **D)** Total proportions of feature intersections for loci, divided by their transcriptional support.

Hairpins with no Rfam hits were then analyzed in the context of their genomic adjacency to other genes. These were subjected to intersection analysis against the NCBI annotations of protein-coding genes (PCGs) and a high-quality annotation of transposable elements (TEs) in fungi [44], where available (Figure 3B). Loci were considered to interact with a feature if their distance is less than 100 nucleotides, additionally labeling loci located entirely within a PCG intron. We found across species that many of the confirmed/no-sequencing-evidence loci arise from PCGs (Figure 3B). The intersection of mi/milRNAs with PCGs is similar to what may be found in animals [49], reflected in high-confidence miRNAs in *Drmel* and *Caenoharbditis elegans* (*Caele*) (Figure 3B). Very rarely, mi/milRNAs from fungi were located entirely within an intron of protein coding genes as one might expect from a mirtron as seen in animals [50] (Figure 3B). In the case of *Drmel* high-confidence miRNAs, around half of those intersecting PCGs were found within introns, while in *Caele* no high-confidence miRNA was found within introns (Figure 3B). Nearly all fungal loci intersections with PCGs were found in exonic regions, with *Cryptococcus neoformans* (*Crneo*) as the primary exception, showing approximately half of its mi/milRNAs within introns (Figure 3B). While this is intriguing in the context of splice-machinery-derived miRNAs [45], more evidence is needed to confirm splicing as the manner of processing of these *Crneo* mi/milRNAs. Mapping loci by their distance from a feature shows several species with near associations to loci that do not intersect (e.g. *Botryosphaeria dothidea* – *Bodot*), possibly pointing to a relationship that may not be directly transcriptionally linked (Figure 3C). TE intersections with mi/milRNA are a likely source of erroneous annotations, as these are known sources of sRNAs (other than miRNAs) associated with genome protection in fungi [1]. Comparing the feature intersections of loci in regard to their confirmation status, we find that a much lower proportion of confirmed loci intersect with TEs (Figure 3D), likely a sign of incorrect annotation.

We found 393 mi/milRNA loci that did not intersect other genomic features (TE, PCG) and also failed to find a homolog in the Rfam dataset (Figure 3B). This number was usually proportional to the number of loci reported for a given species. Considering that we cannot identify any genic features from which these loci derive, we termed these as intergenic loci which were further used for analyzing locus conservation.

### Intergenic fungal mi/milRNAs are conserved in related species

Conservation is strong evidence of the importance of a genomic element. Retention between species points toward purifying selection to maintain a sequence or structure. In plants and animals many miRNA loci are highly ancestral and conserved in sequence and function (e.g. *let-7* in animals, miR166 in plants) [22,52].

To define mi/milRNAs that are conserved between fungal species, we searched for orthologous mi/milRNAs, looking for evidence of a retained hairpin. A simple search for the MAS from a mi/milRNA over sequencing libraries did not reveal any conserved sequences, leading us to explore whether hairpin sequences might be conserved. A genomic search for hairpin sequences is challenging, as large amounts of sequence variation can occur in many regions as long as they maintain the same structure (i.e. compensatory variation) e.g. plant miR156, (Figure S7). To search for conserved hairpin sequences with high-sensitivity, we used HMMER [53]. Assemblies of related fungi, plants, and animals were used as subjects, with hairpins regarded provisionally as homologs following a cutoff of > 0.4 bits / base pair, attempting to normalize for shorter hairpins as is found in animals. As an input, we used only intergenic hairpins (confirmed or no sequencing evidence, Figure 3C) since this reduces the risk of homology due to genic elements not involved with the mi/milRNA. This has a cost of reduced sensitivity, as mi/milRNAs may originate from other genic elements. Figure 4A shows sub-clades of an 18s rRNA tree [51] for a set of related fungal genomes, indicating species that have published mi/milRNAs. Homologs are primarily found only in closely related species based on the strict cutoffs mentioned, with only four examples of homology between class-level designations (*Scscl* mi/milRNAs to *Alternaria brassicicola* – *Albra, Gimar*, and *Bodot*).

**Figure 4.**
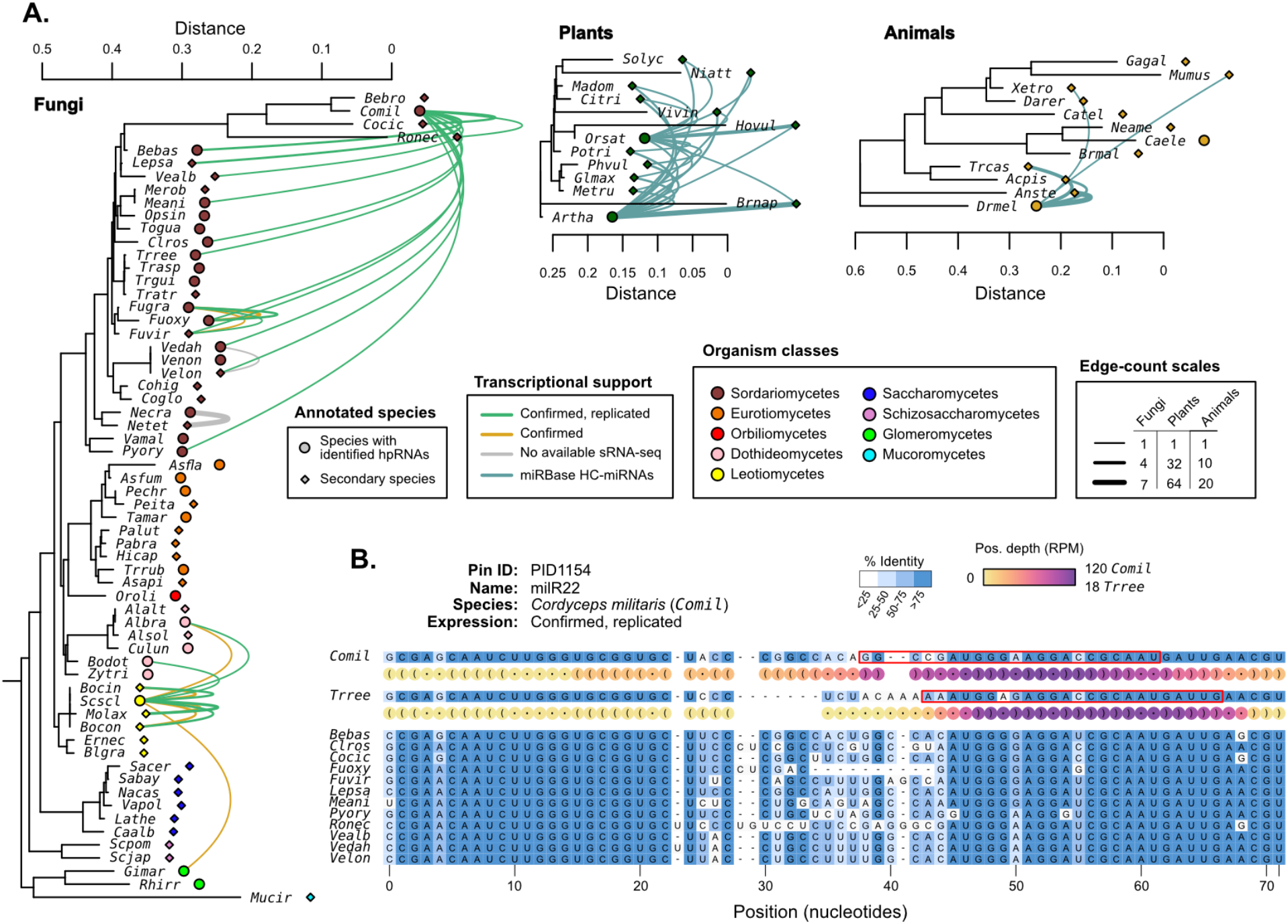
Hairpin RNAs were compared between species to identify possible homologous genes. **A)** 18s rRNA trees [51] (MUSCLE alignment, RAxML maximum likelihood tree) showing species with published mi/milRNA loci (circles) and closely related species with available genomes (diamonds). Connecting edges indicate mi/milRNA loci with highly conserved hairpin sequences in other species (nhmmer, > 0.4 bits / basepair). Edge colors indicate the degree of transcriptional support for the source loci in the context of Figure 2: green – confirmed and replicated loci, yellow – loci confirmed in a single library, grey – loci with no available sequencing libraries. HC-miRNA loci from *Arabidopsis thaliana* (*Artha*) and *Oryza sativa* (*Orsat*) are shown in blue. Edge line width is scaled to indicate the number of connections between two species, with separate scales for each clade. Edge line width indicates the number of connecting edges between two species. **B)** Example of a conserved locus from *Cordyceps militaris* (*Comil*) with other Sordariomycetes. Bases are colored by percent identity. The hairpin structure for *Comil* and *Trichoderma reesei* (*Trree*) are shown in dot-bracket format with colored circles showing scaled depth (RPM) for all positions across available libraries. Most-abundant sequence for these species are highlighted with a red box.

The most highly conserved sequence is milR22 (PID1154) from *Cordyceps militaris* (*Comil*), which is found in 14 Sordariomycetes in our assembly set (Figure 4B). The conservation pattern of milR22 shows a similar structure to that of known miRNAs (Figure S7), pointing to selection for the duplex region. Expression profiling in *Comil* shows that reads primarily come from one hairpin arm, consistent with a hairpin-derived sRNA locus. Profiling was also performed in the three species with putative orthologs and available sequencing data (*Bebas, Fusarium oxysporum*, and *Trichoderma reesei - Trree*). *Trree* was found to produce sRNAs from this putative hairpin with a similar profile to *Comil* (Figure 4B) supporting that these may be orthologous genes. A closer examination of the expression profile for *Comil-*milR22 also shows very high rates of reads that only partially map to the published hairpin (“out-of-bounds”, File S1). This is found in several published mi/milRNA loci, indicating a need for a larger reassessment of locus annotated in these publications.

While there is conservation in fungal mi/milRNAs, it does not compare with the magnitude in plants, where miRNA conservation is widespread (Figure 4A). Animal miRNAs also show signs of conservation, though distinctly lower than plants, possibly a result of their shorter hairpin lengths (Figure 4A). These loci with genomic conservation are strong candidates for biologically important mi/milRNAs, as some selection pressure is plays a part in their retention.

### Well-supported fungal mi/milRNAs have similar characteristics to plant and animal miRNAs

Each prior analyses give different facets of support to published mi/milRNA loci. Merging these diverse forms of evidence can help us categorize and rank this support, as shown in Figure 5. Loci were organized into four progressive tiers of support, with tier 1 being the best supported (Table S5). Tier 1 hairpins are intergenic, found to be conserved between genomes, and are supported by our re-analysis of source sRNA-seq data. Tier 2 loci remove the requirement for genomic conservation and include PCG-derived loci. Tier 3 loci have been confirmed by sRNA-seq data, but come from genomic regions that are likely false positives for hairpins, considering they have inherent secondary structures (Rfam) or are known to produce dsRNA-derived sRNAs (TEs). Tier 4 are likely false positive identifications, as these have critical failings in their sRNA-seq-derived evidence or are positively identified as a structural RNA or TE.

**Figure 5.**
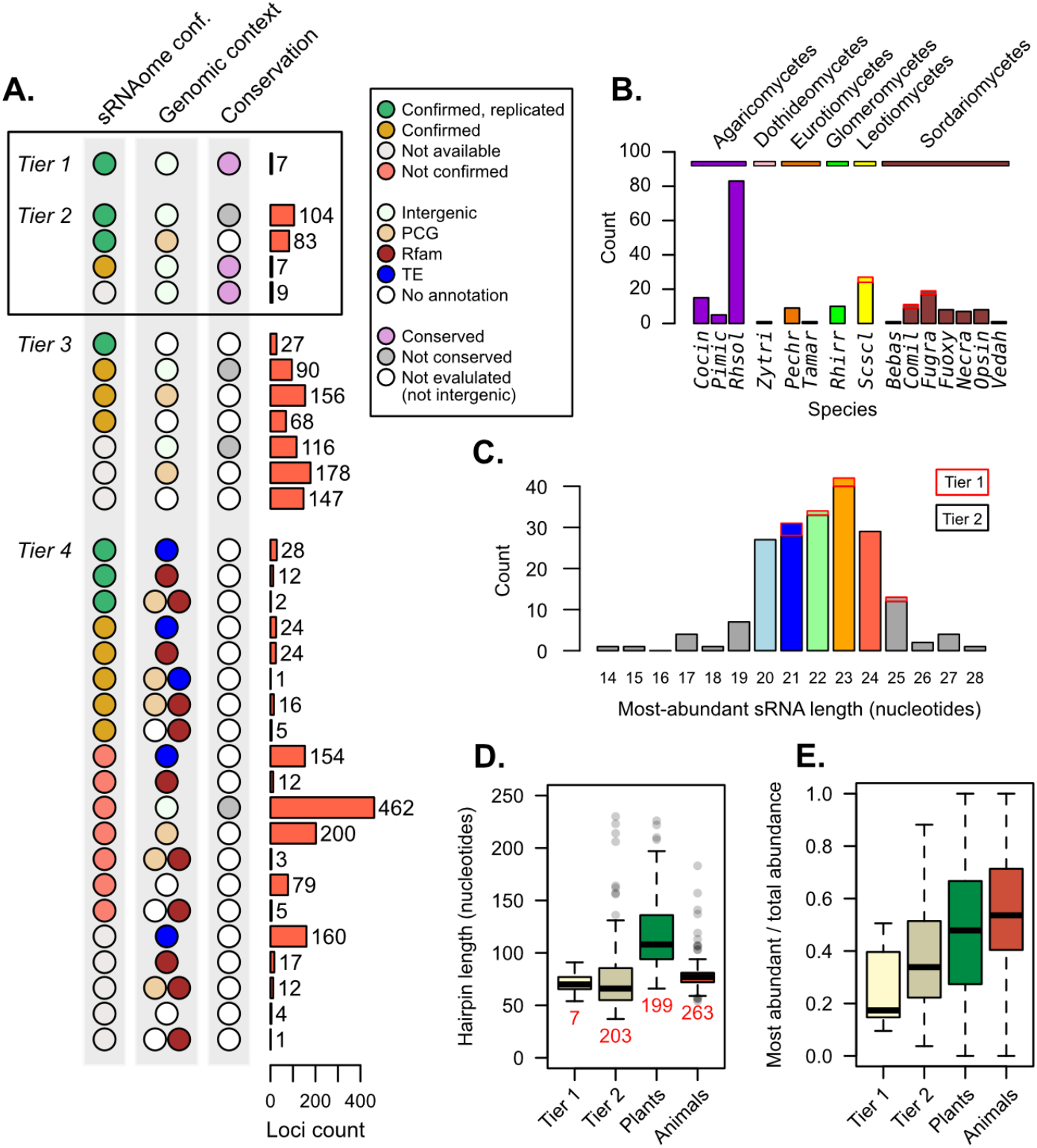
Supporting evidence was used to categorize hairpin RNAs into tier classifications. **A)** Tier classifications of loci based on evidence quality. Colors indicate the support for a locus in the context of figures 2, 3, and 4, with the bars showing the count of loci. Tiers 1 and 2 were analyzed further, looking at **B)** their species (showing loci count) and **C)** the length of the most-abundant sRNA for a locus. For B and C, bar outlines indicate tier of loci (red – tier 1, black – tier 2). **D)** Hairpin lengths of Tier 1 and Tier 2 mi/milRNAs and of HC-miRNAs for plants and animals from miRbase. Length is defined as the duplex-to-duplex distance (including the duplex). **E)** Specificity of sRNA dicing for the same groups, measured by the ratio of the most-abundant sRNA to all sRNAs matching the hairpin strand.

Loci falling into tiers 1 and 2 were considered to have strong support as mi/milRNA loci and were compared to miRNAs in other organisms. 210 loci met this distinction, coming from 15 different fungi from diverse clades (Figure 5B). The most commonly identified species was *Rhizoctonia solani* (*Rhsol*), which had the highest quantity of confirmed and replicated loci of all species (Figure 2E). These top-tier loci tend to fall into similar size ranges as found in plants and animals (Figure 5C). Cleavage by dicer proteins is known to be the causative factor in sRNA size and only rare evidence exists for mi/milRNAs with pathways alternative to dicer [28]. Considering this, loci falling into more atypical ranges (shorter than 19 or longer than 25 nucleotides) may be false positives for mi/milRNA as they might not be indicative of a dicer-derived locus. This is further evidence of a need of standards defining sRNA-profiles of mi/milRNAs in fungi. Hairpin lengths tend to be short in fungal loci, much more similar to those of animals than plants (Figure 5D). Indeed, the inter-duplex length for loci is usually minimally sized, and less than 20 nucleotides in all. A very basic measure of the specificity of processing for a sRNA locus is the proportion of total locus abundance that is represented by the MAS. Here, we show that MAS of top-tier mi/milRNAs in fungi comprise a much lower proportion of locus reads (Figure 5E). This is possibly a sign of differences in the specificity of dicing in fungi or that these loci are not genuine miRNAs or milRNAs. Overall, these characteristics point to similar features to known miRNAs, which is further evidence of an ancient mechanism for miRNAs in eukaryotes.

Using the combined evidence presented in this work, we provide a centralized source of mi/milRNA annotations in fungi. This includes annotations outlined in GFF format (File S2) and complete tables describing extensive details of the loci with supporting evidence for all loci (Table S5). These form a valuable community resource for those looking to reference these loci, with all required details of their genomic coordinates, reference genome, sequence, sRNA profiles, expression, and confidence.

## Discussion

The growing number of publications focused on mi/milRNAs in fungi reflect an increasing awareness of the role these hpRNAs play. Despite the wealth of loci identified in these publications, it is evident that there are persistent problems with filtering out false identifications. Our work finds that only a fraction of these loci remains when performing independent validation. Particularly concerning are those loci failing minimal standards using their own sequencing data, pointing to a serious problem with the standards for locus identification based on sRNA-seq alignment profiles.

It is crucial to adopt a pipeline which is fully auditable for the publication of new sRNA loci, such as a miRNA. This means publishing the unabridged output of any tools used in annotation, including important details such as the sRNA-profile of the locus itself (File S1). This is especially important if sequencing libraries for the study are not made available. Custom analyses and curation should be largely avoided, but if they are used, they must be explained thoroughly and demonstrate supporting evidence found with common tools. Many of the publications citing mi/milRNAs in fungi failed to share even the most basic features of the locus they have identified, sometimes only giving a single sRNA sequence. For an annotation to be of use to the scientific community, full details of the locus should be given, including precursor coordinates, sequences, and predicted folding structures. Plant and animal miRNAs also suffered from this lack of detail, which was a major motivation for establishing of the miRbase repository [54]. Even with complete reporting, low-quality annotations have been pervasive in miRNA annotations [16,38], showing that this challenge is not unique to fungi. Indeed, evidence in this work points to many fungal mi/milRNAs that have qualities similar to low-confidence miRNAs (Figure 2C), further emphasizing a need for stricter standards for annotating these loci. We found that only a small number of loci passed all of the rules laid out for miRNAs by some of the more strict rule-sets (Figure 3D) [37,41]. It is unclear if this is truly representative of the number of loci that might be defined by miRNA rules or if technical challenges are preventing their discovery. An approach more tuned to this clade may be required to assess the question more broadly.

Many tools exist for sRNA-annotation, with most focusing on miRNAs in animals and plants [35–37]. In fungi, at least one tool has been developed for milRNA identification [55], but uses a genome free approach and doesn’t address other categories of loci. The lack of reporting quality and differences in methods between species and studies shows a need for systematic analyses in fungal sRNA loci as has been performed for plants [56] to more clearly grasp the actual types of loci produced in this clade.

Dimensions of a sRNA-locus can tell us a great deal about its synthetic pathway and likely function in an organism. Confirmed hairpins in fungi are much more similar to animals in length – they are very short with little to no terminal loop structure. Phylogenetic comparisons of dicer proteins in eukaryotes show fungal proteins differing vastly from plants and animals where they are much more similar [57]. Animals have a two-step mechanism for dicing hairpins, utilizing two different RNAse III proteins (DCR and Drosha) [47].

However, no evidence points to such a compartmentalization of dicing roles in fungi, further confounding the observed hairpin similarities. Lengths of the MASs from mi/milRNAs broadly fall into expected size ranges for eukaryotes (20-24 nucleotides). Specific DCRs are the causative factor for determining sRNA length [10], but fungi have an average of only two DCR homologs in a species [29] and are thought to be partially-redundant in some cases [28]. This limited radiation of DCR proteins challenges how this range of lengths might be possible. Our results point to low-specificity in fungal mi/milRNA dicing, which might explain the wide range of sizes observed, though this may also be a sign of clade-specific neofunctionalization of DCRs or other unknown protein factors.

Important questions still remain about how these loci may function, target, and affect the expression of target genes. These results are challenging to obtain and frequently require follow-up studies to confirm functionality and role of a proposed mi/milRNA [58]. Evidence of cleavage found in degradome and 5’-RACE experiments hints that fungal mi/milRNAs may function more similarly to plants [31,59–65], as this evidence indicates directed cleavage of a target RNA. However, much more information is needed to develop a detailed view of these dynamics, especially in terms of base-pairing requirements [66] or distinct mechanisms of different AGOs [11]. Our analyses did identify numerous mi/milRNAs which are likely biologically relevant and these may be strong candidates for future analyses of expression, biogenesis, and biological function/role. It is also important to note that validation of sRNA function does not guarantee that it is derived from a mi/milRNA, as there are numerous RDR-derived classes that can likely function similarly. The deeper question that surrounds these elements is whether mi/milRNAs in fungi function in a manner consistent with miRNAs in other organisms, or whether there is any functional distinction between these sRNAs.

The genomic context of fungal mi/milRNAs remains uncertain. While many loci were considered lower quality given their homology with TEs or structural RNAs, these still might be legitimate sources of mi/milRNA. Indeed, some work has concluded that miRNAs [67] may be derived from rRNAs. However, RDR-derived sRNAs are more frequently associated with rRNAs [68], including the qiRNA class in fungi [13]. Finding strong evidence supporting these loci would represent a unique path for biogenesis of mi/milRNAs and could give great insight into the genomic evolution of fungi [69]. These annotations should be treated with caution, as all expressed loci produce short RNAs as a natural result of degradation. Accordingly, these require more attention to conclude that they are actually derived from a DCR protein and are a sRNA locus at all. Considering the enriched proportion of TEs in loci which did not pass confirmation (Figure 3D), loci discovered with these features are likely a symptom false annotation. This is also true for those derived from protein coding regions, though this phenomenon is much more documented [70].

Conservation of mi/milRNAs is a strong sign of their importance. Retention of a processed hairpin between species can be an indicator of selection for retention of function [22]. However, even in systems with widespread conservation, many identified miRNAs are lineage specific. These non-conserved miRNAs are often imprecisely processed and lowly expressed [71] making them frequently designated “low-confidence”. Many fungal mi/milRNAs presented here have these similar characteristics, with nearly all of top-tier loci shown as highly lineage-specific, based on our conservation analysis. Considering that the vast majority of species analyzed here are pathogens, this could be related to adaptation to hosts and/or a function as *trans*-species sRNAs [6]. These sRNAs quickly differentiate in parasites, with evidence of evolution influenced by the host. [23]. The most conserved locus from our work (milR22, *Comil*) is an insect pathogen [72], hinting at a possible *trans*-species role. In all, much more evidence is needed to conclude that these genomically conserved mi/milRNAs are similarly expressed, processed, functional, and facilitate a similar role. Identifying the conservation of these aspects will be an important milestone in the exploration of mi/milRNAs in fungi.

## Conclusion

A large number of sRNAs derived from hairpin sequences have been reported as mi/milRNAs throughout the literature. Here we provide a complete and centralized annotation of these loci, giving all essential data for their exploration. This work identifies that many of the loci fail in basic validation of the locus structure and expression, highlighting a need for better standards of annotation in fungal mi/milRNA loci. Around 10% of loci were found to fall in the highest tiers of support, with these validations and annotations provided as a community resource. We found that these loci frequently arise from coding regions of the genome, including some from introns in genes. Genomically conserved mi/milRNAs also provide insight into the possible retention of these loci and their function. Overall, we see that fungal mi/milRNAs are similar in many ways to those of animals and plants though it is not always clear to which they are more, varying in aspects of sRNA length, hairpin length, and mode of action. These results provide one of the first clade-wide explorations of this topic and allow greater perspective into the still-developing topics of mi/milRNAs in fungi.

## Methods and Materials

### Finding published fungal mi/milRNAs

To assess the state of mi/milRNAs in fungi, we searched for publications related to “miRNAs” and “miRNA-like RNAs” in fungal species. We performed this search on the Pubmed web interface using the following term: (“Fungi”[Text Word] OR “Fungus”[Text Word]) AND (“miRNA” OR “milRNA”) NOT Review[Publication Type], yielding 148 results (July 29 2022). These were subsequently filtered by hand as to whether they referenced these types of sRNAs in a fungal species. Publications were then assessed for whether they identify sequences or loci related to mi/milRNA, extracting all relevant sequence and coordinate information. Only discoveries based on small-RNA-sequencing were considered. Other information extracted was the accession numbers associated with sequencing repositories, any indications about the replication of a given mi/milRNA in multiple libraries, and the genome assembly used to identify loci. When assemblies were incompletely identified, the correct genome was inferred by manually matching coordinates (where available) to likely assemblies.

Support for mi/milRNA genes was analyzed in two contexts: genetic evidence supporting their synthesis pathway and functional evidence supporting targeting relationships. Confirmed genetic evidence constitutes studies that used knock-out, knock-downs, or over-expression to identify genetic dependencies for at least one mi/milRNA. For proving function of a mi/milRNA, studies that relied only on target prediction or indirect perturbations (i.e. a knockout of essential sRNA genes) were considered weak evidence. Molecular evidence of cleavage (5’-RACE, degradome) or observing the effects of direct modulation of a specific mi/milRNA abundance (i.e. through target mimics or sRNA supplementation) were considered strong evidence.

A complete record of loci was produced based on the available data, including the coordinates, chromosome/scaffold (with NCBI identifiers), strand, hairpin sequence, genome/assembly, and the published mature sRNA-sequences. Where coordinate information was lacking but a hairpin sequence was available, coordinates were inferred using blastn [73]. Hairpins were folded using RNAfold (default options) [40]. The duplex was estimated based on the pairing for the published mature sequence, including a conventional 2 nucleotide 5’ overhang on the mature and star sequences. To be considered valid, a hairpin must 1) contain the reported MAS, 2) have no secondary structures present in the duplex region, and 3) have no more than 20 base positions in the duplex that do not have a paired sequence.

### Recovering incompletely reported loci

Loci which only reported a mature sequence or coordinates were submitted to a pipeline to recover a candidate hairpin sequence and genomic source. The pipeline first identified a likely locality for the hairpin. For those with mature coordinates, this was used as center of the locality. For those with only a single mature sequence provided, genome alignment of this sequence with bowtie [74] was used to find candidate localities. For those with two mature sequences, genome alignments of these two sequences were intersected so that a locality must contain both. In the case of multiple localities found, were tested in an arbitrary order (random). Testing involved dividing a locality into sequential candidate hairpins, 10 each of sizes 150, 300, and 600 nucleotides, making 30 candidates in all. These were folded and assessed for validity as shown prior. The recovered hairpin was chosen based on the most frequently identified duplex sequence among the hairpins. The chosen hairpin passing these requirements was reported for a mi/milRNA and all subsequent localities were ignored. Precision and sensitivity of the recovery pipeline was tested using the mature sequences only for loci with valid and fully reported hairpin sequences, including miRbase miRNA loci for *Artha, Orsat, Drmel*, and *Caele*.

### Expression profiling of hairpins

For those studies with publicly available data, libraries were downloaded and processed. *Artha* control libraries were used from a prior publication of the lead author (PRJNA543296 – stem tissue) [23]. *Drmel* control libraries were chosen from a recent submission from NCBI-SRA (PRJNA636660 – whole body). Adapters were identified from a set of commonly used Illumina and sRNA-seq adapters. Trimming was performed with cutadapt (-a [adapter_seq] –minimum-length 10 –maximum-length 50 –overlap 4 –max-n 0) [75]. ShortStack [41] was used in alignment-only mode to perform a weighted bowtie alignment [74] to the assembly. This approach includes alignments for most multi-mapping reads, choosing a single placement based on the local abundance of uniquely mapping reads [34]. Trim and alignment rates were confirmed to be mostly similar against reported rates in source publications, where available. All RPM calculations in this work were performed using all genomically-aligned reads as the denominator.

Hairpins were then assessed according to rules from other publications and tools relating to miRNA identification [16,34,35,37]. The rules outlined in these works are summarized in Table S6, including a basic hairpin rule set introduced by this work. These rules were tested on the expression profiles of small RNAs aligned to the hairpins, which do not exceed the bounds of the hairpin sequences. In some cases, the mature sequence was not the MAS in the locus.

### Homology to structural RNAs

To determine if loci are derived from a structural RNA, we used the Rfam database [48]. First, we filtered a set of Rfam entries looking for families which contain a reference to at least one of the fungi with published mi/milRNA (Table S5). For the figures, families were generalized to categories and any annotations of small RNA loci were not considered as a structural RNA. Using this filtered list, we performed a blastn [73] search for every hairpin sequence, considering hits with a bitscore >= 50.

### Genomic adjacency

To find if loci are derived or near to protein coding genes or transposable elements, we relied on annotations of these features. Assemblies and associated gene annotations for this step were all obtained from NCBI genbank accessions (with a GCA prefix). Gene annotations were filtered to only protein coding genes (“gene_biotype=protein_coding”). TE annotations were provided from [44]. When the assembly associated with a mi/milRNA did not have an NCBI entry, or failed to have one or both of the annotations, we chose the most complete assembly that did. We used blastn alignment of the hairpin sequence to translate between genomes where necessary. Genomic distance was determined using bedtools closest [76], reporting the closest feature to the hairpin locus on either strand. This was repeated for TE and protein coding gene annotations. In the case of a tie, a single closest feature is reported, with Tes favored in the case of a tie in distance (or intersect). Distances of 100 nucleotides or less were considered intersections in terms of calling intergenic loci.

### Finding orthologous hairpin sequences

To identify putative orthologs of mi/milRNA in other fungi, we utilized genomic sequence searches. Subject genomes included all species with published mi/milRNAs shown in this study. In addition, we also included many close and distant relative species with annotated assemblies available in NCBI, especially in clades where the prior assemblies had limited resolution. Sequence searches were performed using hmmer [53]. Databases from assemblies were formed with the command makehmmerdb –sa_freq 2. All intergenic hairpins were then searched against the databases using the command nhmmer –tblout. Search output was filtered to include only orthologous hits (excluding hits in the same species) and redundant hits in a species, retaining only the best candidate by bitscore. To control for length of hairpins, positive hits were filtered to have a bitscore / input sequence length > 0.4. Profiles for milR22 homologs in other species were performed by aligning all libraries of said species to their hairpin profile, using bowtie [74].

### Building phylogenetic trees

To compare organismal taxonomy, phylogenetic trees were constructed based on 18s ribosomal data from the SILVA database (v138.1) [51]. Single sequences for each species were obtained from the reference NR99 aligned dataset when available and taken from the partial database when not available there. Alignments were performed to remove gaps and align the partial sequences to the NR99 alignments using MUSCLE (3.8.31) with default settings [77]. Maximum likelihood trees were produced for each organismal kingdom separately (fungi, plants, and animals) using raxml -m GTRCAT -p 9182 [78].

## List of supplementals

### Supplementary Tables

Table S1 – List of publications reporting mi/milRNA from fungi referred in this work. We show the publication identifier used in this work, the organism it pertains to, the publication title, journal, URL, whether the miRNA/milRNA reported were obtained by sRNA-seq analysis (Y:yes; N:no), details of why a study was excluded, whether sRNA-seq data is available (Y: yes, N:no), whether mi/milRNA loci coordinates are provided (Y:yes, N:no), whether biological replication of results has been performed (Y:yes, N:no), whether genetic evidence is provided (Y:yes; N:no), type of targeting evidence provided (prediction, degradome, 5’ RACE, *in vitro* support for targeting, and detection of target gene modulation through indirect approaches knock-outs of biosynthetic genes and direct approaches which modify the expression of the functional sRNA), and bioinformatic tool used for mi/milRNA identification. This also gives information on the native assembly used for the primary identification of the mi/milRNA loci cited in a publication, including assembly sources, IDs, names, and URLs where applicable.

Table S2 – List of species considered in this work. We show species abbreviations (abbv), full-names (species), taxon identifiers (taxid), secondary-names (synonyms), kingdom, phylum, class, order, family, genus, whether they are a species reported mi/milRNAs analyzed in this work (1: yes; 0: no), relevance to humans, hosts for pathogens/parasites, and corresponding genome assemblies used for genomic context and homolog identification (ncbi_assembly).

Table S3 – sRNA-seq libraries used for re-assessment of reported mi/milRNA loci. We show the publication citations used in this work, SRR accession of the libraries, adapter sequence used for trimming, read length of the library, number of reads in the raw library (raw_depth), and the percentage of raw reads in the following categories: trimmed with adapters (have_adap), reads passing □utadapt filters (pass_filter), uniquely mapping reads, multi-mapping reads, percent non-mapping reads, and aligned reads resulting from unique and multi-mapper placement (aligned). Where available, this also gives the alignment rates given by source publications and a manually curated note.

Table S4 – Rules testing data for miRNAs and fungal libraries. We show identifiers for the mi/milRNA locus, citation, and source sRNA-seq library. Locus abundance, hairpin strand abundance, and percent stranding are shown. Other features include the abundance and sequence for the most-abundant sequence determined in this work (MAS), the inferred star for this MAS (MAS*), and the MAS cited in the source publication. Locus and cited MAS designation shown in accordance with Figure 2A. Rule passing is shown as a string, with the rule citation shown with a two-letter prefix (from Table S6), with the subsequent string pointing to rules passing (1) and failing (0) in numerical order.

Table S5 – Table defining the confirmation status for a mi/milRNA for all reported loci. We show identifiers for the locus, publication, native and NCBI assemblies used in its analysis, hairpin coordinates used for each assembly, and the most abundant sequence, indicating whether it has been corrected by our sequencing re-analysis. Locus details like the hairpin sequence, folding, minimum free energy, and boundary coordinates of the duplex within the hairpin are provided. We report identification details, including the level of coordinate and locus detail provided by the source publication and a locus’s validation status. Confirmation details include the rule-set confirmation status, interactions with Rfam (including Rfam descriptions), intersections with genomic features, the intergenomic conservation status, and the tier classification of all reported loci. Relevant figures are indicated where available for data columns.

Table S6 – miRNA annotation rules used in this work. Rule identifiers use the prefixes “ax” [37], “ku [35], “mb” [16], and “ss” [41]. General rule category and more detailed rule-descriptions are presented for each rule.

### Supplementary Figures

Figure S1 – Analysis of locus count by citation. We show the number of loci reported by species in each publication. Abbreviations are provided in Table S2 and publications are provided in Table S1. Colors are respective to their taxonomic class shown in Figure 1B.

Figure S2 – Accuracy for recovery pipeline. A) Cumulative density function of the number of map locations for mature mi/milRNAs sequences in their corresponding genome assemblies. Sensitivity is displayed for the cutoff of 10 mature mi/milRNA alignments in the genome. Precision is modeled based on a single mature mi/milRNA alignment, randomly chosen from possible alignments. B) Actual precision and sensitivity values computed for loci with a known source location as positive. Recoveries are done using only mature sequences for loci with known loci.

Figure S3 – Evaluation of reported mi/milRNA loci using published rules for plant and animal miRNAs (description of rules is provided in Table S6). Plants-LC: set of low-confidence miRNAs from miRbase for plants (Viridiplantae set); Plants-HC: set of high-confidence miRNAs from miRbase for plants (Viridiplantae set); Animals-LC: set of low-confidence miRNAs from miRbase for animals (Metazoa set); Animals-HC: set of high-confidence miRNAs from miRbase for animals (Metazoa set).

Figure S4 – Cumulative density function of raw library depths for source libraries reporting mi/milRNAs. We show the proportion of libraries with a given raw read depth. 94.1% of the libraries are 5 million reads or more in depth.

Figure S5 – Loci count in the context of their confirmation by sRNA-seq. A) Proportion and loci count for loci reported from each citation, B) the annotation tool/approach, and C) the length of the most-abundant sequence. Colors represent loci which are invalid (dark grey), failed to confirm by sRNA-seq (red), have no available sRNA-seq data (light grey), are confirmed in one library (yellow) or are confirmed in multiple libraries.

Figure S6 – Repetitiveness of unfiltered loci in related genomes. A) Cumulative density function of the number of blast hits in included genomes for hairpins from fungal mi/milRNAs. Top hits are labeled with the target genome species identifier and the mi/milRNA which has a blast hit. B) Expanded view of the count of species where bba-milR4 is found, separated by kingdom.

Figure S7 – Multiple sequence alignment for homologs of miR159 across plant species. Positions are colored by their column-wise percent identity. Fold and MASs are shown for *Artha*.

### Supplementary Files

File S1 – Text file of the sRNA abundance profiles for all mi/milRNA with available sRNA-seq data. Here is given the locus and library identifiers, abundance of the locus and all sRNAs aligned to the hairpin strand (l=length, a=abundance). Most-abundant sequence (MAS) and cited MASs are shown for each locus. Lower case nucleotides indicate mismatches with the reference.

File S2 – Annotations of mi/milRNAs (gffs). Includes annotations for mi/milRNAs in native genomes (where they were first annotated) and best-blast-hits (bbh) in NCBI-sourced assemblies. Annotations also contain two versions, one with all mi/milRNA loci and a second with only top tier loci (tiers 1 and 2, denoted “best”).

## Contributions

NRJ, EAV, and JMA were responsible for the conception and overall structure of the project. NRJ performed the annotation, recovery, and all analyses. NRJ wrote the principal manuscript, with substantial revisions from EAV and JMA. LFL gave important contributions to the direction of the project and final revisions of the manuscript. The authors have declared no competing interests.

## Acknowledgements

José David Fernandez and Evelyn Sánchez (Centro de Genómica y Bioinformática, Universidad Mayor) and Dr. Matthew Hasenjager (National Institute for Mathematical and Biological Synthesis, University of Tennessee, Knoxville) for help in accessing source papers and supplemental data. Dr. Anna Muszewska (Institute of Biochemistry and Biophysics, Polish Academy of Sciences) for providing the complete set of TE annotations referenced in their publication [44]. Dr. Michael Frietag (Center for Genome Research and Biocomputing, Oregon State University) for providing the provisional genome sequences used in their publication [28]. Dr. Yulong Wang (Anhui Agricultural University) for providing details into the methodology for their publication [72].

