## Supplementary material for "Comprehensive re-analysis of hairpin RNAs in fungi reveals ancestral links": Figures S1-7

**Figure S1**

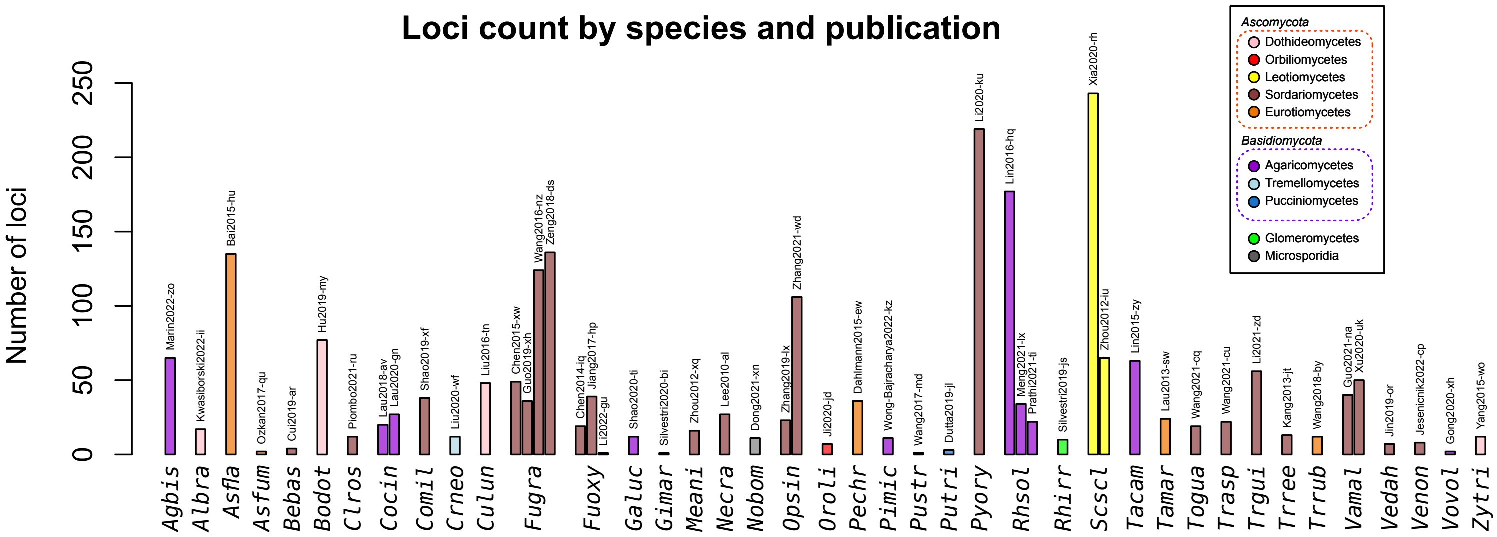

Loci count by species and publication
Number of loci
Ascomycota
Dothideomycetes
Orbiliomycetes
Leotiomycetes
Sordariomycetes
Eurotiomycetes
Basidiomycota
Agaricomycetes
Tremellomycetes
Pucciniomycetes
Glomeromycetes
Microsporidia
Agbis
Albra
Asfla
Asfum
Bebas
Bodot
Clros
Cocin
Comil
Crneo
Culun
Fugra
Fuoxy
Galuc
Gimar
Meani
Necra
Nobom
Opsin
Oroli
Pechr
Pimic
Pustr
Putri
Pyory
Rhsol
Rhirr
Scscl
Tacam
Tamar
Togua
Trasp
Trgui
Trree
Trrub
Vamal
Vedah
Venon
Vovol
Zytri

Figure S2

A.

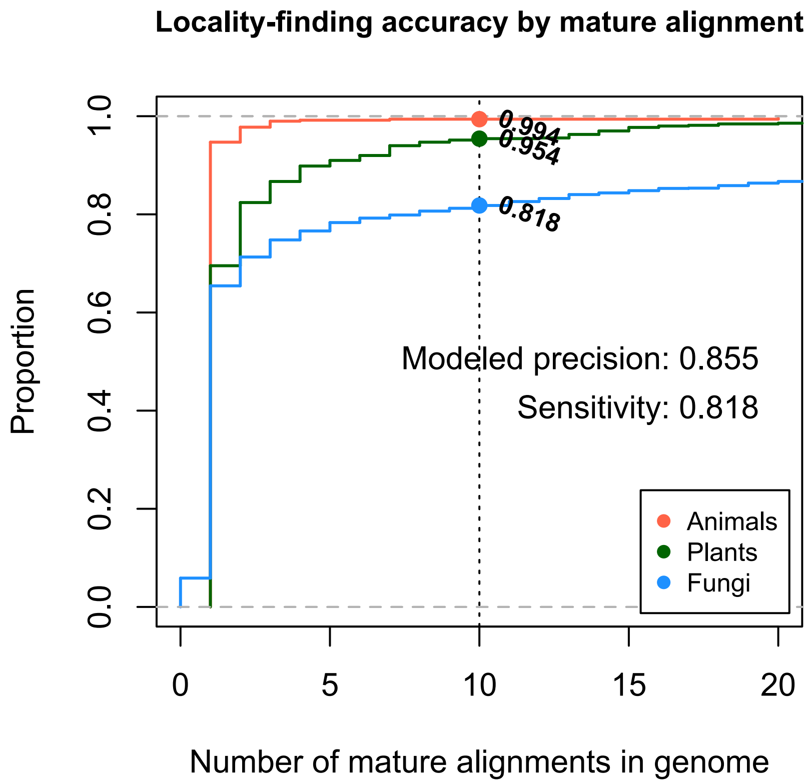
Locality-finding accuracy by mature alignment
Proportion
0.0
0.2
0.4
0.6
0.8
1.0
0
5
10
15
20
Number of mature alignments in genome
0.994
0.954
0.818
Modeled precision: 0.855
Sensitivity: 0.818
Animals
Plants
Fungi

B.

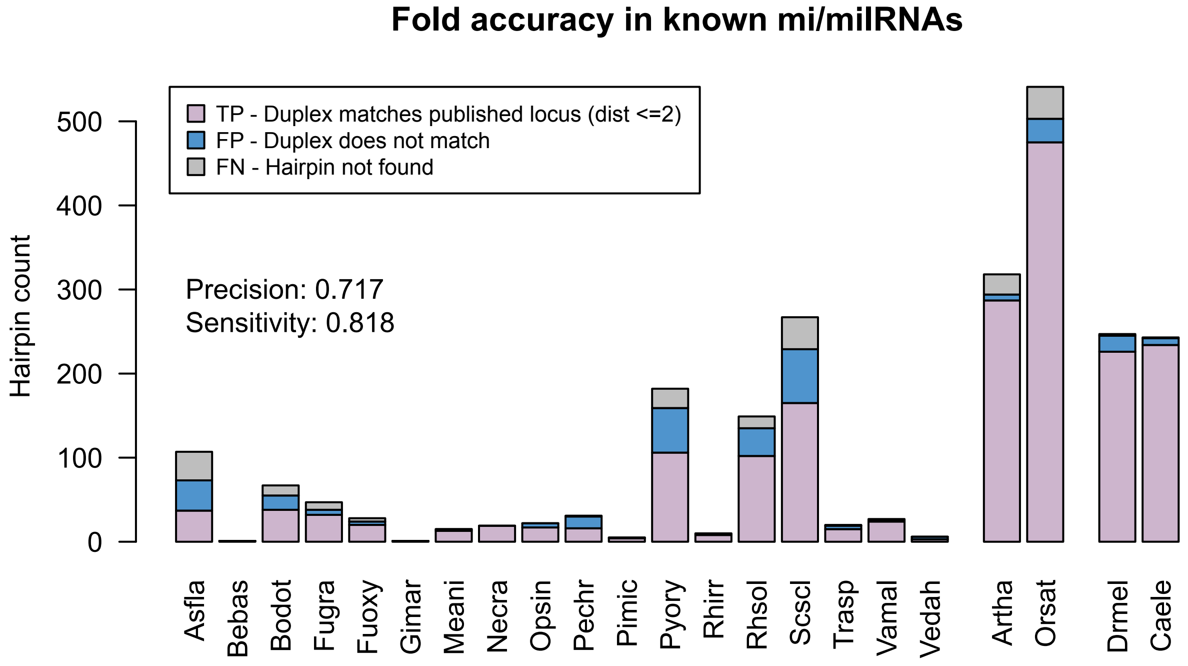
Fold accuracy in known mi/milRNAs
TP - Duplex matches published locus (dist <=2)
FP - Duplex does not match
FN - Hairpin not found
Hairpin count
0
100
200
300
400
500
Precision: 0.717
Sensitivity: 0.818
Asfla
Bebas
Bodot
Fugra
Fuoxy
Gimar
Meani
Necra
Opsin
Pechr
Pimic
Pyory
Rhirr
Rhsol
Scscl
Trasp
Vamal
Vedah
Artha
Orsat
Drmel
Caele

**Figure S3**

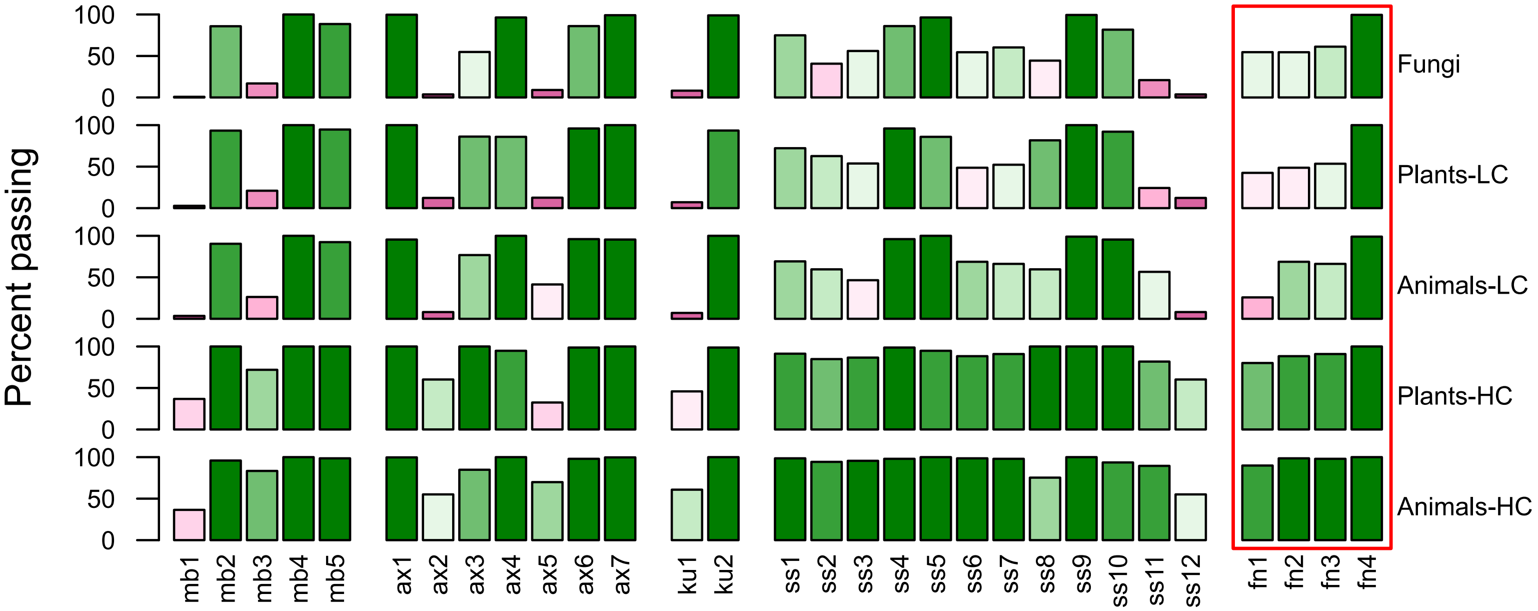
Percent passing
100
50
0
Fungi
Plants-LC
Animals-LC
Plants-HC
Animals-HC
mb1 mb2 mb3 mb4 mb5
ax1 ax2 ax3 ax4 ax5 ax6 ax7
ku1 ku2
ss1 ss2 ss3 ss4 ss5 ss6 ss7 ss8 ss9 ss10 ss11 ss12
fn1 fn2 fn3 fn4

**Figure S4**

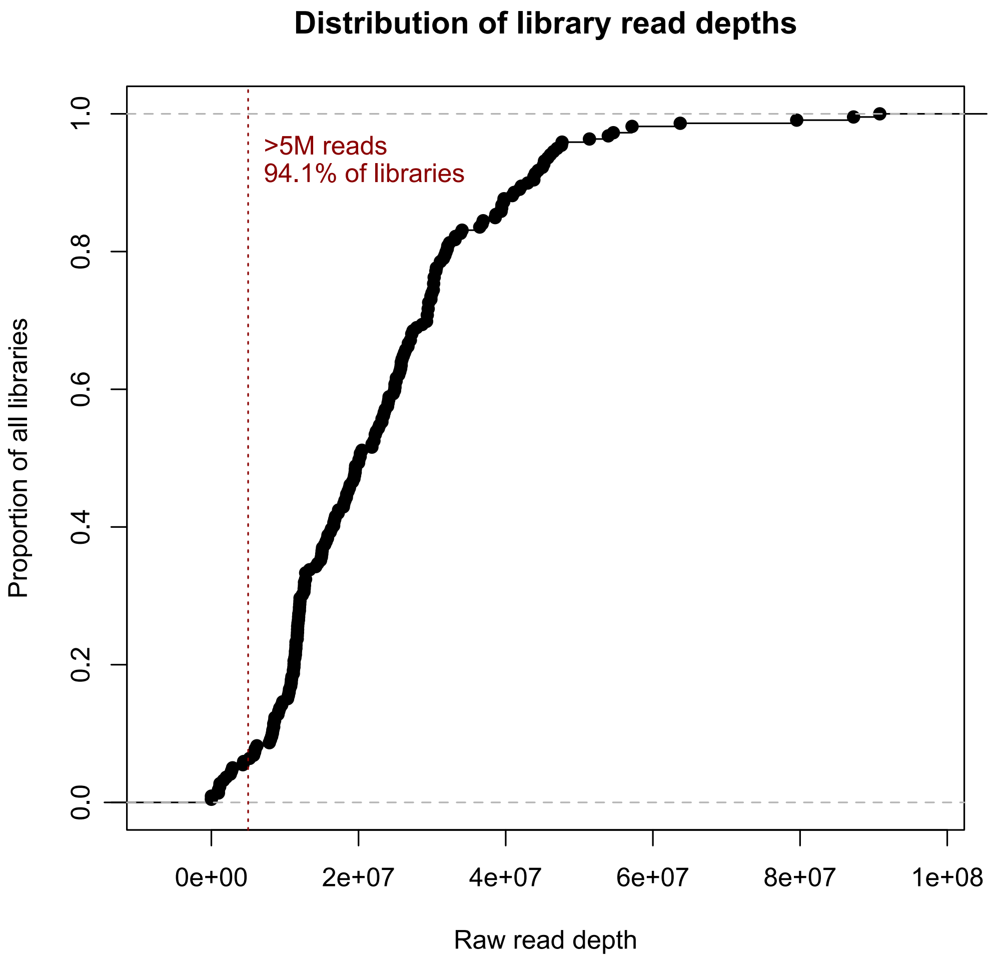

Distribution of library read depths
>5M reads
94.1% of libraries
Proportion of all libraries
0.0
0.2
0.4
0.6
0.8
1.0
0e+00
2e+07
4e+07
6e+07
8e+07
1e+08
Raw read depth

#### Figure S5

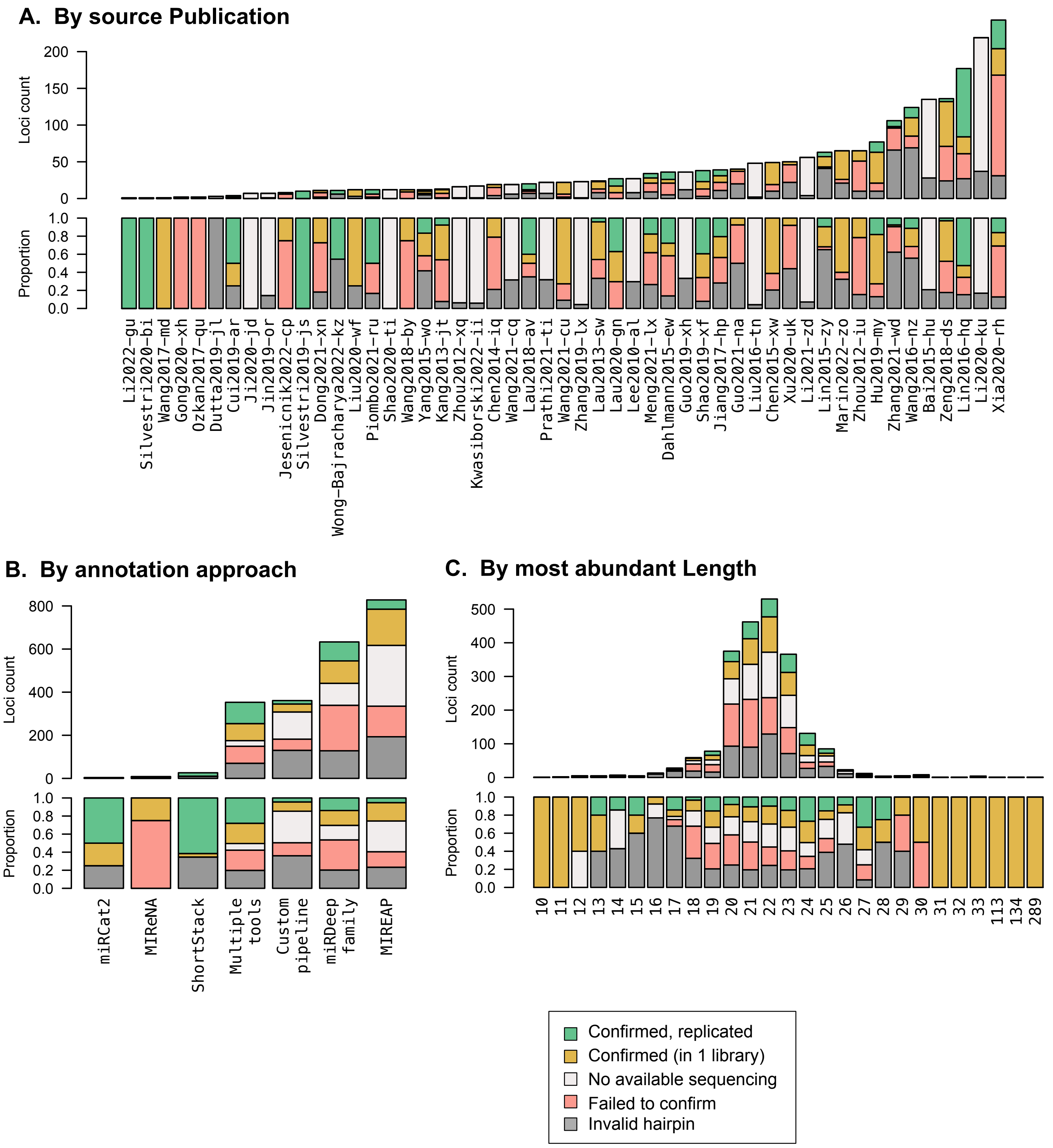

A. By source Publication
Loci count
Proportion
Li2022-gu
Silvestri2020-bi
Wang2017-md
Gong2020-xh
Ozkan2017-qu
Dutta2019-jl
Cui2019-ar
Ji2020-jd
Jin2019-or
Jesenicnik2022-cp
Silvestri2019-js
Dong2021-xn
Wong-Bajracharya2022-kz
Liu2020-wf
Piombo2021-ru
Shao2020-ti
Wang2018-by
Yang2015-wo
Kang2013-jt
Zhou2012-xq
Kwasiborski2022-ii
Chen2014-iq
Wang2021-cq
Lau2018-av
Prathi2021-ti
Wang2021-cu
Zhang2019-lx
Lau2013-sw
Lau2020-gn
Lee2010-al
Meng2021-lx
Dahlmann2015-ew
Guo2019-xh
Shao2019-xf
Jiang2017-hp
Guo2021-na
Liu2016-tn
Chen2015-xw
Xu2020-uk
Li2021-zd
Lin2015-zy
Marin2022-zo
Zhou2012-iu
Hu2019-my
Zhang2021-wd
Wang2016-nz
Bai2015-hu
Zeng2018-ds
Lin2016-hq
Li2020-ku
Xia2020-rh
B. By annotation approach
Loci count
Proportion
miRCat2
MIReNA
ShortStack
Multiple tools
Custom pipeline
miRDeep family
MIREAP
C. By most abundant Length
Loci count
Proportion
113
134
289
Confirmed, replicated
Confirmed (in 1 library)
No available sequencing
Failed to confirm
Invalid hairpin

#### Figure S6

**A.** Blast hits of mi/milRNAs within other genomes

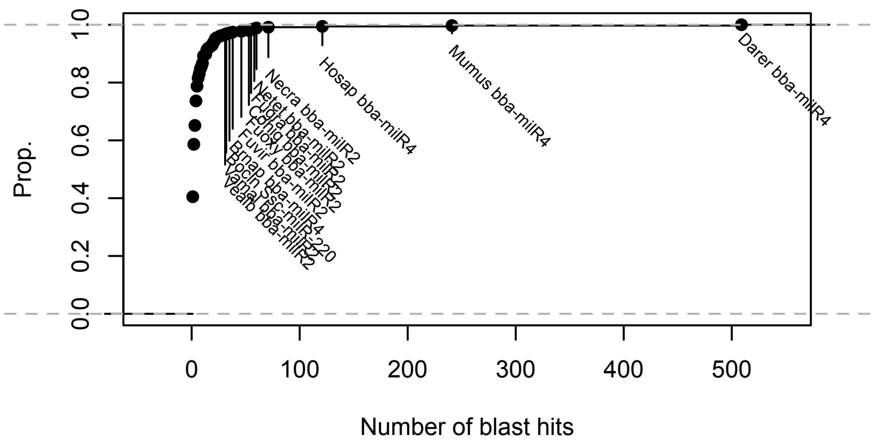

**B.** Frequency of genomic occurences for *bba*-milR4 within other species

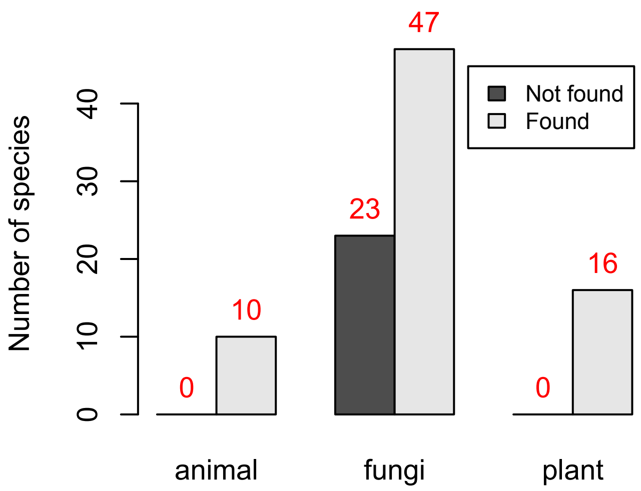

### Figure S7

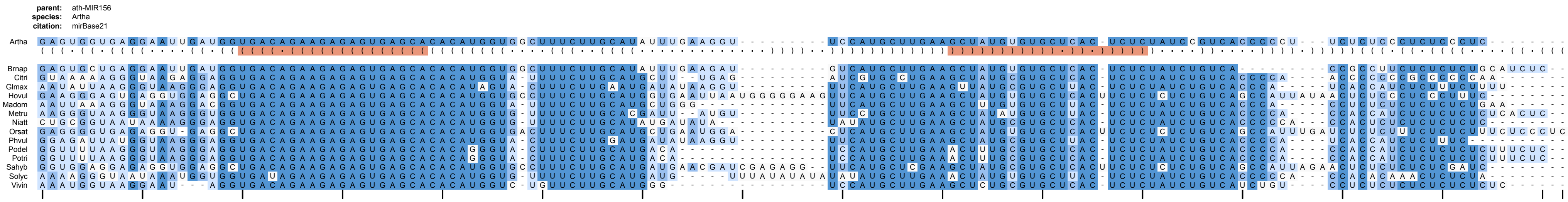
parent: ath-MIR156
species: Artha
citation: mirBase21
Artha
Brnap
Citri
Glmax
Hovul
Madom
Metru
Niatt
Orsat
Phvul
Podel
Potri
Sahyb
Solyc
Vivin
